# SummArIzeR: Simplifying cross-database enrichment result clustering and annotation via large language models

**DOI:** 10.1101/2025.05.28.656331

**Authors:** Marie Brinkmann, Michael Bonelli, Anela Tosevska

## Abstract

**Motivation:** Enrichment analysis across multiple databases often results in a high level of redundancy due to overlapping terms, complicating the interpretation of biological data. To address this, we developed SummArIzeR, an R package to cluster and annotate enrichment results across multiple databases, enabling fast, intuitive interpretation and comparison across multiple conditions. SummArIzeR clusters enrichment results based on shared genes, calculates a pooled p-value for each cluster and facilitates the cluster annotation using large-language models. It further allows an easyly interpretable vizualisisation of the results.

**Results:** Compared to existing tools, SummArIzeR provides unbiased and fast cluster annotation using large language models. We demonstrate that SummArIzeR achieves clustering comparable to manual curation while offering superior grouping based on shared underlying genes.

**Availability and Implementation:** The SummArIzeR package is available as an open-source R package, with a comprehensive user manual provided in its GitHub repository: https://github.com/bonellilab/SummArIzeR.

## Introduction

Multi-OMICs techniques are powerful tools that allow the exploration of multiple biological layers, from the genome to the transcriptome and epigenome (1,2). These techniques generate large amounts of data that, after correct analysis and interpretation, can give meaningful insights into biological mechanisms and their perturbations. Despite the multitude of existing methods for analysis of such data, the interpretation remains a significant challenge, especially when more layers of information are integrated. Gene-set databases, such as **Gene Ontology (GO)** and **MSigDB**, have helped remarkably by enabling gene-set enrichment analysis and linking gene-sets to biological contexts (3,4).

In order to understand biological mechanisms, it’s often valuable to combine enrichment results from multiple databases. Yet, this can introduce redundancy, with many overlapping or similar terms, which can create inflated and complex visualization.

A variety of methods and tools have been developed to address this issue (**Figure 1**). Traditionally, manually clustering and annotating these terms requires expertise and intuition and relies on term semantic similarity rather than shared underlying genes, which can often lead to misleading interpretations. While flexible as it is not limited to specific databases, this method is time-consuming, requires substantial biological expertise, and does not consider the underlying gene sets associated with each term.

**Figure 1.**
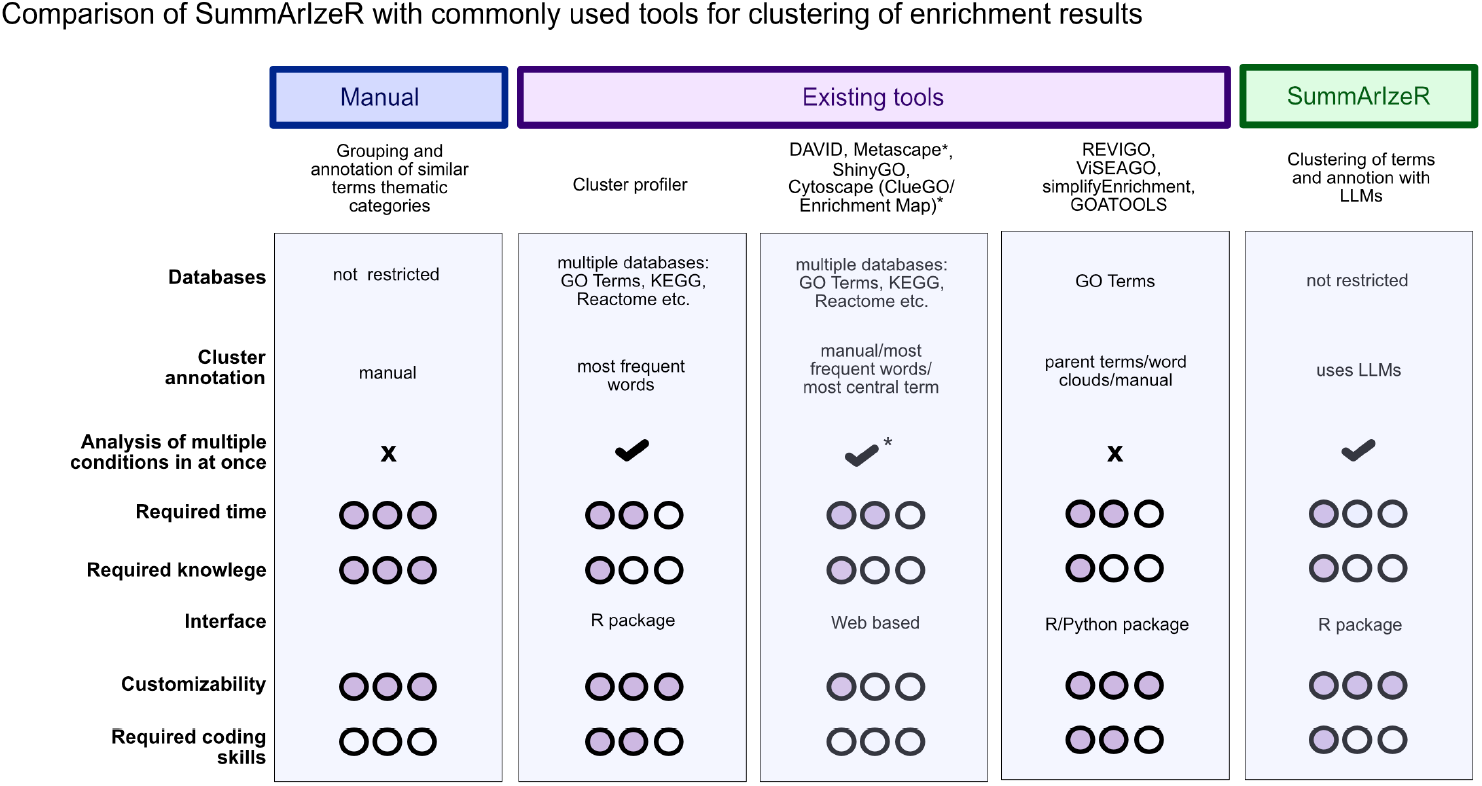
Comparison of SummArIzeR with existing tools for clustering enrichment results.

Several clustering-based solutions, such as **REVIGO, VISEAGO, simplifyEnrichment**, and **GOATOOLS**, improve on this by grouping similar GO terms based on term semantic similarity. However, these tools are generally restricted to Gene Ontology terms and can only process one condition at a time (5–8).

To extend to multiple databases and conditions, web-based tools like **DAVID, Metascape, ShinyGO**, and **Cytoscape** offer user-friendly interfaces. Although these platforms are easy accessible, the analysis is time-consuming and their output figures are often limited in customization (9–12). For users comfortable with coding, **clusterProfiler** offers an R-based solution with more flexibility, but it requires a certain level of programming experience (13).

While all these tools address clustering to some degree, annotation of the resulting clusters remains limited. Most rely on parent term relationships or word-cloud-based algorithms, both of which can lose important biological information (**Figure 1**) (5–13).

The rise of large language models (LLMs) like ChatGPT offers new possibilities in bioinformatics. LLMs have already begun supporting NGS and multi-OMICs data analysis, such as assisting in cell-type annotation for single-cell sequencing datasets (14–16). They now present an exciting opportunity: unbiased, fast, and biologically informed annotation of multiple enrichment terms.

Here we introduce SummArIzeR, an R package designed to streamline enrichment analysis across multiple conditions and databases. SummArIzeR clusters enriched terms based on shared genes and uses LLMs such as ChatGPT to generate meaningful cluster annotations, allowing to investigate biological insights of OMICs-data. We believe that SummArIzeR overcomes the afore-mentioned limitations. It enables simultaneous analysis of multiple conditions, such as different cell types, disease states, or treatment groups and supports the separate evaluation of up- and downregulated genes (**Figures 1)**. With a simple R interface requiring only minimal coding knowledge, users can quickly perform analyses.

## Methods

SummArIzeR was developed using R 4.2.2, utilizing the **Enrichr** (17**), dplyr, tidyr, stringr**, and **igraph** packages. Output plots are based on **Complex Heatmap** (18) and ggplot2. Additional plots were generated using **ggVennDiagram**.

### Methods within SummArIzeR

#### Enrichment analysis

Input gene lists are processed to extract gene identifiers and, optionally, log2 fold-change values. Functional enrichment is performed using the **Enrichr** Application Programming Interface (API), and results are filtered based on an adjusted p-value threshold of <0.05 and a minimum number of genes per term. When specified, genes are separated into upregulated and downregulated subsets based on a user-defined log2 fold-change threshold.

To support the analysis of multiple experimental conditions simultaneously, the enrichment pipeline is applied across combinations of user-defined experimental groupings and selected **Enrichr** databases. Enrichment results from all conditions and databases are aggregated, with a configurable filter applied to retain the top-ranking terms per condition based on statistical significance.

#### Clustering of terms

Identified terms are clustered based on methodologies previously implemented in the **simplifyEnrichment** package (6). Briefly, a binary gene-term incidence matrix is constructed, to calculate pairwise Jaccard distances. A weighted undirected graph is built, and low-weight edges are pruned based on a user-defined threshold. Clusters are defined by Community Walktrap algorithm (**igraph** package), allowing assignment of biologically coherent clusters. In brief, the walktrap community detection is based on random walks through a network in order to measure distance between vertices, where a random walk gets trapped in denser neighbourhoods of the network that correspon to communities (19).

#### Cluster annotation and p-Value pooling

Generated clusters are annotated, by creating a promt for any LLM. P-values for terms within clusters are pooled using a choice of four different methods: Fisher’s method (20), Stouffer’s method (21), a weighted Z-test (22) or the Cauchy combination test (23). By default, pooled p-values are capped at a minimum threshold for visualisation purposes.

Clusters get annotated by mapping manual summaries to cluster IDs. For each cluster and condition, p-values are pooled using the selected method. If regulation status (up/down) is available, it is retained. Redundant columns are removed, and unique term and gene counts are calculated per cluster.

#### Visualization: Heatmaps and Bubble Plots

Enrichment results can be visualized using heatmaps and bubble plots. Heatmaps show −log10(pooled p-values), with optional bar plot annotations for term counts per cluster. Furthermore, conditions and clusters can be further grouped using hierarchical clustering. Bubble plots displayes condition vs. cluster annotation, with bubble size representing significance and color reflecting the number of genes enriched in the specific cluster.

### Example data

To demonstrate the usage of SummArIzeR and compare it to published results we used published data from Kugler et al. (24). Data was processed as described in Kugler et al. and used as an input for SummArIzeR.

## Results

### SummArIzeR Workflow

In a first step, users can upload a table including gene names and condition names **(Figure 2A)**. For a separate analysis of up-and downregulated genes, a column indicating log2-fold change has to be present. The user can select any database available from the extensive Enrichr library (17) and perform the enrichment analysis within SummArIzeR. The clustering can be influenced by setting a treshold parameter. An interactive igraph network vizualises the term clustering **(Figure 2B)**, and a threshold evaluation plot can be used to check modularity, connected cluster and cluster count for different tresholds **(Figure 2C)**. A unique feature of SummArIzeR is its use of LLMs to annotate clusters based on the underlying terms, preserving individual term information in an efficient and unbiased way **(Figures 1, 2A)**. The output is a table listing clusters, their annotations, associated terms, and underlying genes. This avoids a “black-box” approach, empowering users to investigate clustering decisions and customize their own visualizations. Graphical outputs include heatmaps and bubble plots (ggplot2 object), enabling straightforward comparison across multiple conditions **(Figure 1D)**.

**Figure 2.**
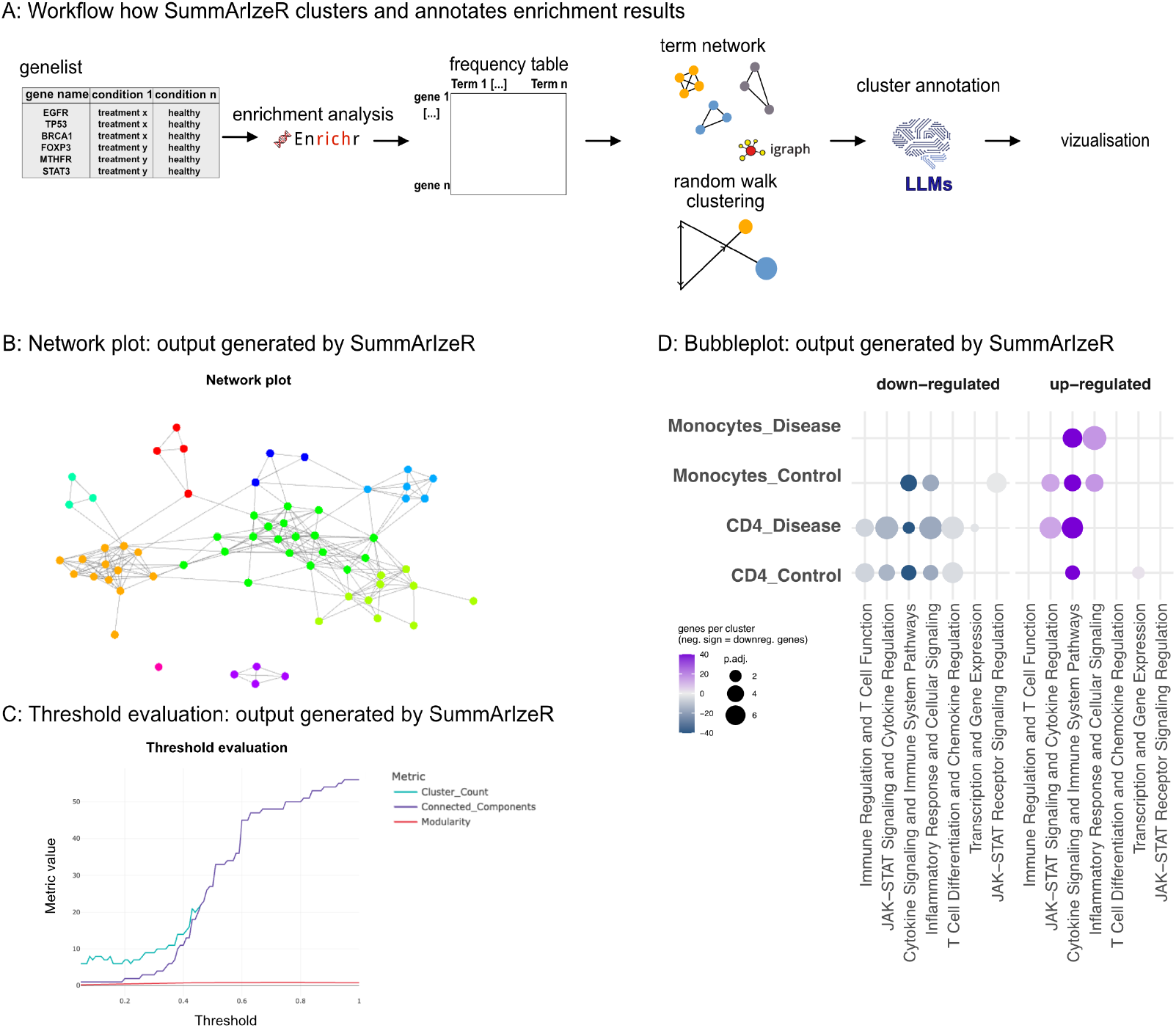
**A**. Core workflow of SummArIzeR **B**. Example of a network plot (SummArIzeR output) with colors indicating cluster membership. **C**. Threshold evaluation plot (SummArIzeR output) showing how cluster count, number of connected components, and modularity change with varying thresholds. **D**. Bubble plot (SummArIzeR output) generated with synthetic data, using the databases: GO_Biological_Process_2023, Reactome_2022, and BioPlanet_2019.

### Dataset: Cytokine stimulation of fibroblast-like synoviocytes (FLSs) induces unique and shared transcriptional programmes

To demonstrate how SummArIzeR improves the visualization and interpretation of enrichment results, we applied it to a previously published dataset by our group that required comparison across multiple conditions and was affected by high term redundancy, requiring manual annotation (24). The dataset contains bulk-RNA sequencing results of fibroblast-like synoviocytes, stimulated with different cytokines. We performed the enrichment analysis using the “GO-Biological-function_2021” database, selecting the top five terms per condition, comparable to the analysis performed in the publication (Kugler et al., Figure 2). In the original analysis, terms were manually clustered into 6 groups based on term semantic similarity. With SummArIzeR, we performed the enrichment analysis under the same conditions. We observed similar cluster distribution, however, some differences could be observed in the term assignments (**Figure 3A, Figure 3B**).

**Figure 3.**
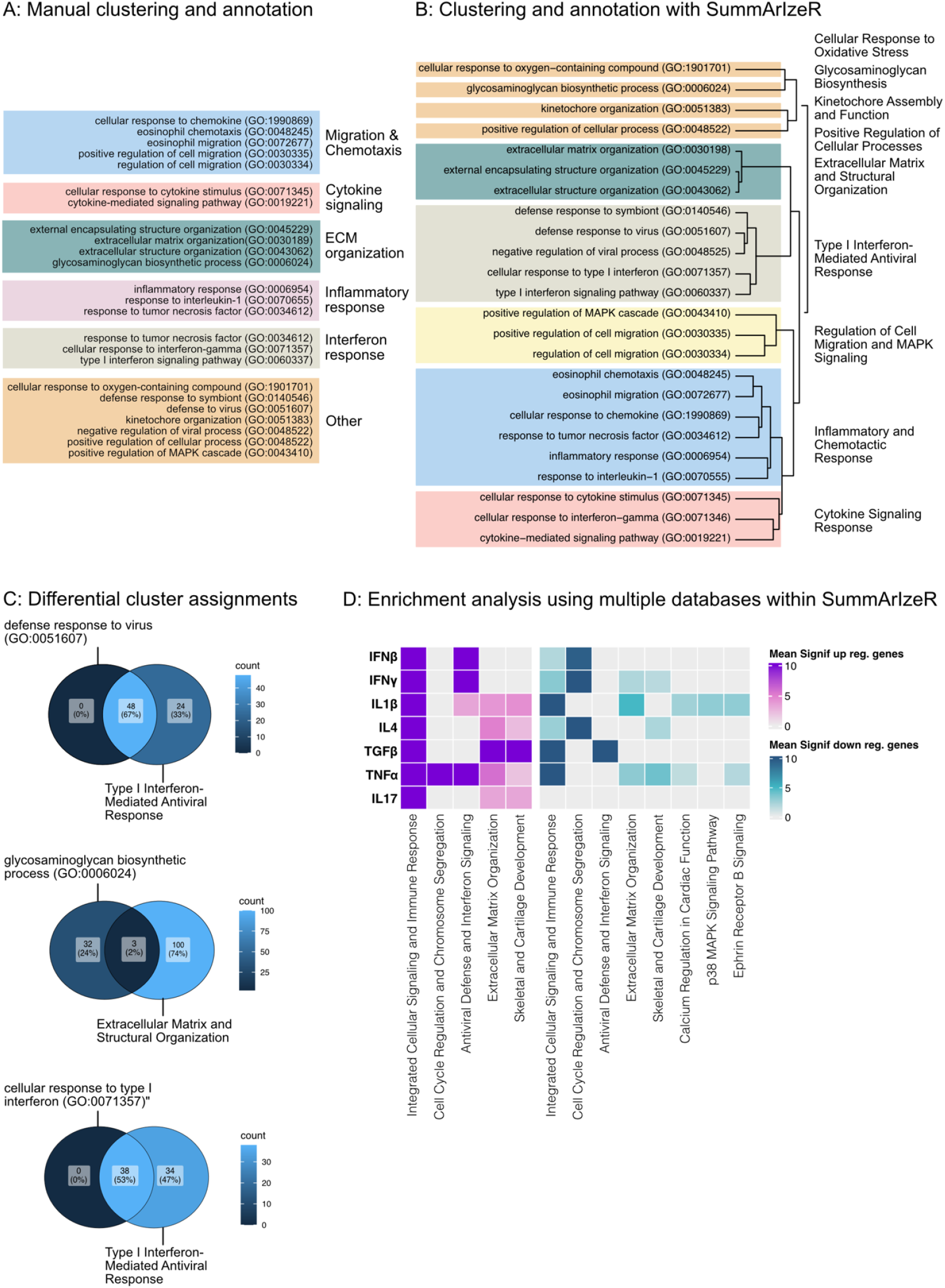
**A**. Manual clustering of GO terms, following the approach used in Kugler et al., Figure 2: The top five enriched terms per condition (based on the GO_Biological_Process_2021 database) were manually grouped. **B**. Automated clustering using SummArIzeR with the same database (GO_Biological_Process_2021) and top five hits per condition. **C**. Examples of differing cluster compositions, illustrated with Venn diagrams showing gene overlap between terms and clusters generated with SummArIzeR, visualized as color gradients. **D**. Re-analysis of the dataset using SummArIzeR with different databases (GO_Biological_Process_2025 and BioPlanet_2019), based on the top five hits per condition (Heatmap visualization).

The term “defense response to virus (GO:0051607)” was not assigned to a group in the original analysis. (**Figure 3A**), but it was assigned to the “Type I Interferone Mediated Antiviral Response” cluster by SummArIzeR. The enrichment analysis showed, that 48 genes in the dataset matched the term. All of these genes overlapped with enriched genes found in the cluster “Type I Interferone Mediated Antiviral Response” after excluding the term “defense response to virus (GO:0051607)” from this cluster (**Figure 3C**).

The term “glycosaminoglycan biosynthetic process (GO:0006024)” was manually clustered in the group “ECM organization” (**Figure 3A**). In contrast, SummArIzeR clustered the term seperatly. Investigating overlapping genes revealed, that only 2% of the genes overlapped with the cluster “Extracellular Matrix and Structural Organization”, after excluding the term “glycosaminoglycan biosynthetic process (GO:0006024)” from the cluster (**Figure 3C**).

The term “cellular response to type I interferon (GO:0071357)” was manually annotated in a group called “Interferone response”, together with the term “cellular response to interferone-gamma (GO:0070655)” (**Figure 3A**). The enrichment analysis showed, that 38 genes in the dataset matched the term. All of these genes overlapped with enriched genes found in the cluster “Type I Interferone Mediated Antiviral Response” after excluding the term “cellular response to type I interferon (GO:0071357)” (**Figure 3C**).

As a next step, using SummArIzeR, we could extend the enrichment analyses to include other databases and visualize them (**Figure 3D**). We used the “GO_Biological_Process_2025” and “BioPlanet_2019” databases, selecting the top 5 hits for each condition. We performed the cluster annotation with ChatGPT-4 and could further show, that other LLMs like DeepSeek, Claude, Perplexity AI and Gemini AI led to similar cluster annotation (**Supplementary Data**). Including additional databases uncovered additional insights, such as the involvement of cell cycle-related genes, cartilage development pathways, calcium regulation mechanisms, and ephrin receptor B signaling - processes relevant to fibroblast activation in synovial tissue. Importantly, separating upregulated and downregulated genes revealed more nuanced patterns; for example, TGF-β stimulation led to both upregulation and downregulation of genes involved in cellular signaling and immune responses, highlighting the complex regulatory dynamics.

These results could clearly demonstrate that SummArIzeR is a powerful tool for enrichment analysis, enabling systematic, unbiased clustering and annotation of enriched terms. It supports efficient visualization and direct comparison of multiple conditions, facilitating accurate and comprehensive data interpretation.

## Discussion

To date, no tool has provided an easy and systematic approach for clustering and annotating enrichment results across multiple databases. With SummArIzeR, we introduce a user-friendly R-package that enables grouping of enrichment terms into clusters, annotation of these clusters, and direct comparison of p-values across multiple conditions.

SummArIzeR offers several advantages over manual annotation and existing tools. It provides an unbiased, reliable, and user-friendly approach, generating outputs that support fully customizable visualisation. We compared clustering and annotation results obtained with SummArIzeR to those generated through manual grouping of GO terms. SummArIzeR produced comparable clustering and annotations, while offering additional advantages by grouping terms based on shared underlying genes, even when term names did not directly overlap. Furthermore, SummArIzeR revealed a more comprehensive enrichment result compared to the original analysis. It performed enrichment analysis separately for each condition and independently for upregulated and downregulated genes, ensuring that important results, especially from conditions with lower enrichment signals, are not overlooked.

The main limitation is that SummArIzeR currently relies on **Enrichr** for enrichment analysis, which requires stable internet connection when performing the analysis and does at the moment not allow for user-defined gene sets addition. Additional functionalities to address these limitations are planned for future updates, such as the use of offline enrichment and the ability to add custom gene sets. Careful database selection remains crucial, especially regarding organism-and tissue-specific relevance.

While LLMs facilitate cluster annotation, clustering more than 200 terms may occasionally result in incomplete output. This limitation is expected to improve as more advanced LLM versions become available. However, SummArIzeR’s transparent use of LLM outputs allows users to review and potentially adapt the generated annotation results.

Overall, we believe that SummArIzeR simplifies enrichment analysis across multiple databases and conditions, making it a valuable and practical addition to the R toolkit for sequencing data interpretation.

## Code Availability

The SummArIzeR package (v2) is available as an open-source R package, with a comprehensive user manual provided in its GitHub repository (2). All versions of the software are archived on Zenodo (25). The full source code and example datasets used to reproduce the analyses are openly accessible through the GitHub repository: https://github.com/bonellilab/SummArIzeR.

## Acknowledgements

We thank the Division of Rheumatology at the Medical University of Vienna for critical feedback with using and testing SummArIzeR. We thank Dr. Stefan Badelt for his critically reviewing of the manuscript.

## Authorship contributions

Marie Brinkmann: Investigation, Conceptualization, Methodology, Software, Writing – original draft, Visualization Michael Bonelli: Resources, Supervision, Writing – review and editing Anela Tosevska: Validation, Investigation, Conceptualization, Methodology, Software, Supervision, Project administration

## References

1. Dai X, Shen L. Advances and Trends in Omics Technology Development. Front Med. 2022;9:911861.

2. Hayes CN, Nakahara H, Ono A, Tsuge M, Oka S. From Omics to Multi-Omics: A Review of Advantages and Tradeoffs. Genes. 2024 Nov 29;15(12):1551.

3. The Gene Ontology Consortium. The Gene Ontology Resource: 20 years and still GOing strong. Nucleic Acids Res. 2019 Jan 8;47(D1):D330–8.

4. Dharmesh D. tBhuva, Gordon K. Smyth, Alexandra Garnham. msigdb [Internet]. Bioconductor; [cited 2025 Apr 28]. Available from: https://bioconductor.org/packages/msigdb

5. Klopfenstein DV, Zhang L, Pedersen BS, Ramírez F, Warwick Vesztrocy A, Naldi A, et al. GOATOOLS: A Python library for Gene Ontology analyses. Sci Rep. 2018 Jul 18;8(1):10872.

6. Gu Z, Hübschmann D. simplifyEnrichment: A Bioconductor Package for Clustering and Visualizing Functional Enrichment Results. Genomics Proteomics Bioinformatics. 2023 Feb;21(1):190–202.

7. Brionne A, Juanchich A, Hennequet-Antier C. ViSEAGO: a Bioconductor package for clustering biological functions using Gene Ontology and semantic similarity. BioData Min. 2019 Dec;12(1):16.

8. Supek F, Bošnjak M, Škunca N, Šmuc T. REVIGO summarizes and visualizes long lists of gene ontology terms. PloS One. 2011;6(7):e21800.

9. Bindea G, Mlecnik B, Hackl H, Charoentong P, Tosolini M, Kirilovsky A, et al. ClueGO: a Cytoscape plug-in to decipher functionally grouped gene ontology and pathway annotation networks. Bioinforma Oxf Engl. 2009 Apr 15;25(8):1091–3.

10. Sherman BT, Hao M, Qiu J, Jiao X, Baseler MW, Lane HC, et al. DAVID: a web server for functional enrichment analysis and functional annotation of gene lists (2021 update). Nucleic Acids Res. 2022 Jul 5;50(W1):W216–21.

11. Zhou Y, Zhou B, Pache L, Chang M, Khodabakhshi AH, Tanaseichuk O, et al. Metascape provides a biologist-oriented resource for the analysis of systems-level datasets. Nat Commun. 2019 Apr 3;10(1):1523.

12. Ge SX, Jung D, Yao R. ShinyGO: a graphical gene-set enrichment tool for animals and plants. Valencia A, editor. Bioinformatics. 2020 Apr 15;36(8):2628–9.

13. Xu S, Hu E, Cai Y, Xie Z, Luo X, Zhan L, et al. Using clusterProfiler to characterize multiomics data. Nat Protoc. 2024 Nov;19(11):3292–320.

14. Sarumi OA, Heider D. Large language models and their applications in bioinformatics. Comput Struct Biotechnol J. 2024 Dec;23:3498–505.

15. Hou W, Ji Z. Assessing GPT-4 for cell type annotation in single-cell RNA-seq analysis. Nat Methods. 2024 Aug;21(8):1462–5.

16. Enhancing functional gene set analysis with large language models. Nat Methods. 2025 Jan;22(1):22–3.

17. Xie Z, Bailey A, Kuleshov MV, Clarke DJB, Evangelista JE, Jenkins SL, et al. Gene Set Knowledge Discovery with Enrichr. Curr Protoc. 2021 Mar;1(3):e90.

18. Gu Z, Eils R, Schlesner M. Complex heatmaps reveal patterns and correlations in multidimensional genomic data. Bioinformatics. 2016 Sep 15;32(18):2847–9.

19. Pons P, Latapy M. Computing Communities in Large Networks Using Random Walks. In: Yolum pInar, Güngör T, Gürgen F, Özturan C, editors. Computer and Information Sciences - ISCIS 2005 [Internet]. Berlin, Heidelberg: Springer Berlin Heidelberg; 2005 [cited 2025 May 13]. p. 284–93. (Hutchison D, Kanade T, Kittler J, Kleinberg JM, Mattern F, Mitchell JC, et al., editors. Lecture Notes in Computer Science; vol. 3733). Available from: http://link.springer.com/10.1007/11569596_31

20. Fisher RA.Statistical Methods for Research Workers. In: Kotz S, Johnson NL, editors. Breakthroughs in Statistics [Internet]. New York, NY: Springer New York; 1992 [cited 2025 May 13]. p. 66–70. (Springer Series in Statistics). Available from: http://link.springer.com/10.1007/978-1-4612-4380-9_6

21. Stouffer, S. A., Suchman, E. A., Devinney, L. C., Star, S. A., & Williams, R. M., Jr. The American soldier: Adjustment during army life. (Studies in social psychology in World War II). Princeton Univ. Press. Rinceton Univ Press. 1949;

22. Zaykin DV. Optimally weighted Z-test is a powerful method for combining probabilities in meta-analysis: Optimally weighted Z-test is a powerful method. J Evol Biol. 2011 Aug;24(8):1836–41.

23. Liu Y, Xie J. Cauchy Combination Test: A Powerful Test With Analytic p -Value Calculation Under Arbitrary Dependency Structures. J Am Stat Assoc. 2020 Jan 2;115(529):393–402.

24. Kugler M, Dellinger M, Kartnig F, Müller L, Preglej T, Heinz LX, et al. Cytokine-directed cellular cross-talk imprints synovial pathotypes in rheumatoid arthritis. Ann Rheum Dis. 2023 Sep;82(9):1142–52.

25. Brinkmann M, Tosevska A. SummArIzeR [Internet]. Zenodo; 2025 [cited 2025 May 15]. Available from: https://zenodo.org/doi/10.5281/zenodo.15188400

