## SupplementaryData for "SummArIzeR: Simplifying cross-database enrichment result clustering and annotation via large language models"

**Cluster annotation (Figure 3D), comparing different large language models:**

```
#Integrating LLM annotation

## Summary Terms with ChatGPT 4-turbo

cluster_summary <- c(
  '1' = 'Skeletal and Cartilage Development',
  '2' = 'Integrated Cellular Signaling and Immune Response',
  '3' = 'Cell Cycle Regulation and Chromosome Segregation',
  '4' = 'Antiviral Defense and Interferon Signaling',
  '5' = 'Calcium Regulation in Cardiac Function',
  '6' = 'Extracellular Matrix Organization',
  '7' = 'Ephrin Receptor B Signaling',
  '8' = 'p38 MAPK Signaling Pathway'
)
##Summary Terms with DeepSeek

cluster_summary <- c(
  '1' = 'Skeletal and Muscular System Development',
  '2' = 'Inflammatory and Immune Signaling Pathways',
  '3' = 'Mitotic Cell Cycle and Chromosome Segregation',
  '4' = 'Antiviral Defense and Interferon Signaling',
  '5' = 'Calcium Ion Regulation and Cardiac Muscle Contraction',
  '6' = 'Extracellular Matrix Organization',
  '7' = 'Ephrin Receptor Signaling',
  '8' = 'p38 MAPK Signaling Pathway'
)

##Summary Terms with Claude

cluster_summary <- c(
  '1' = 'Skeletal and Cartilage Development',
  '2' = 'Cytokine Signaling and Inflammatory Response',
  '3' = 'Mitotic Cell Cycle Regulation',
  '4' = 'Antiviral Immune Response',
  '5' = 'Calcium Ion Regulation',
  '6' = 'Extracellular Matrix Organization',
  '7' = 'Ephrin Signaling',
  '8' = 'p38 MAPK Signaling'
)

##Summary Terms with PerplexityAI

cluster_summary <- c(
  '1' = 'Skeletal and Musculoskeletal System Development',
  '2' = 'Immune Response and Intracellular Signaling Regulation',
  '3' = 'Mitotic Cell Cycle and Spindle Assembly Regulation',
  '4' = 'Antiviral Defense and Interferon-Mediated Immune Signaling',
  '5' = 'Regulation of Calcium Signaling and Cardiac Muscle Contraction',
  '6' = 'Extracellular Matrix Organization and Collagen Pathways',
```

```
  '7' = 'Ephrin Receptor Signaling Pathway',
  '8' = 'p38 MAPK Signaling Pathway'
)

#Gemini AI

cluster_summary <- c(
'1' = 'Skeletal and Muscle Development',
'2' = 'Immune and Inflammatory Responses and Signaling',
'3' = 'Mitotic Cell Cycle and Spindle Organization',
'4' = 'Antiviral Defense and Interferon Signaling',
'5' = 'Regulation of Calcium Ion Release in Muscle Contraction',
'6' = 'Extracellular Matrix Organization',
'7' = 'Ephrin Receptor B Signaling',
'8' = 'p38 MAPK Signaling')
```
